# Deeply conserved susceptibility in a multi-host, multi-parasite system

**DOI:** 10.1101/424549

**Authors:** Lisa N. Barrow, Sabrina M. McNew, Nora Mitchell, Spencer C. Galen, Holly L. Lutz, Heather Skeen, Thomas Valqui, Jason D. Weckstein, Christopher C. Witt

## Abstract

Variation in susceptibility is ubiquitous in multi-host, multi-parasite assemblages, and can have profound implications for ecology and evolution. The extent to which susceptibility is phylogenetically conserved among hosts is poorly understood and has rarely been appropriately tested. We screened for haemosporidian parasites in 3983 birds representing 40 families and 523 species, spanning ~4500 meters elevation in the tropical Andes. To quantify the influence of host phylogeny on infection status, we applied Bayesian phylogenetic multilevel models that included a suite of environmental, spatial, temporal, life history, and ecological predictors. We found evidence of deeply-conserved susceptibility across the avian tree; host phylogeny explained substantial variation in infection rate, and results were robust to phylogenetic uncertainty. Our study suggests that susceptibility is governed, in part, by conserved, latent aspects of anti-parasite defense. This demonstrates the importance of deep phylogeny for understanding the outcomes of present-day ecological interactions.

*Statement of authorship:* LNB, SMM, NM, and CCW designed the study; SMM, SCG, HLL, HS, TV, JDW, and CCW collected the data; LNB and NM analyzed the data; LNB, NM, and CCW wrote the paper with input from all authors.

*Data accessibility statement:* Specimen information is available from the Arctos database (arctosdb.org) and in supplementary tables (Appendix S1). Files used for analysis will be archived in Dryad. DOI: XXX.

## INTRODUCTION

Susceptibility to parasites and pathogens can affect the fitness of individuals, the structure of communities, and the evolutionary success of lineages. Therefore, the causes of variation in susceptibility among hosts are of paramount importance to ecology and evolution. Host individuals may vary in susceptibility because of differences in exposure or defense (Gaunt 1995; Barrett *et al*. 2009; Savage *et al*. 2011; Atkinson *et al*. 2013). Host species also vary in susceptibility (Power & Mitchell 2004; Searle *et al*. 2011), and interspecific variation can be explained, in part, by variation in aspects of life history, morphology, environment, and behavior (Scheuerlein & Ricklefs 2004; Garamszegi & Møller 2012; Johnson *et al*. 2012; Lutz *et al*. 2015). Additional interspecific variation may be explained by unique ‘species effects’ (e.g., Pulgarín-R *et al*. 2018; Ricklefs *et al*. 2018), attributable to unique, derived species traits, such as molecular genetic aspects of the immune system (Martin *et al*. 2005; Ellison *et al*. 2015).

The extent to which susceptibility to parasites shows a conserved pattern of evolution across the host phylogeny has seldom been addressed. If susceptibility is conserved, it would indicate that the real-time outcome of an ecological interaction is partly contingent on deep-time evolutionary history. Parasites can affect host biogeography, macroevolution, and community assembly (Holt 1977; van Riper *et al*. 1986; Hatcher *et al*. 2006; Bradley *et al*. 2008, Holt & Bonsall 2017), and it follows that conserved susceptibility could constrain phylogenetic community structure and contribute to conserved rates of speciation, extinction, or secondary sympatry. From a broad perspective, parasite clades tend to have phylogenetic limits to their host ranges. For example, haemosporidian parasite genera tend to infect vertebrate hosts in a single class or subclass (Galen *et al*. 2018). Paired host and parasite clades can form dynamic multi-host, multi-parasite assemblages, with host-parasite linkages proliferating by host-switching (Ricklefs *et al*. 2004; Doña *et al*. 2018; Fecchio *et al*. 2018). Host-switching can occur across large phylogenetic gaps within these host and parasite clades (Anderson 2000; Beadell *et al*. 2009; Ricklefs *et al*. 2014; Suh *et al*. 2016), indicating deeply conserved compatibility. The key question we address in this paper concerns the extent to which host species exhibit phylogenetically conserved patterns of susceptibility within a multi-host, multi-parasite assemblage.

One possibility is that susceptibility is labile rather than phylogenetically conserved across the extent of multi-host, multi-parasite systems. Under this ‘lability hypothesis’, variation in susceptibility would be entirely attributable to host species, populations, and individuals. There are at least four lines of supporting evidence for the lability hypothesis. First, temporal and spatial variation in parasite pressure is profound (Bennett & Cameron 1974; Merilä *et al*. 1995; Svensson-Coelho *et al*. 2013). Second, aspects of host ecological and behavioral niches tend to evolve quickly (Blomberg *et al*. 2003; Schreeg *et al*. 2010; Zhang *et al*. 2017), and these can have substantial effects on exposure to parasite vectors (Garvin & Remsen 1997; Walther *et al*. 1999; Scheuerlein & Ricklefs 2004). Third, simple regulatory or structural genetic changes in the immune system can increase or eliminate susceptibility over short time-scales, as suggested by rapid changes in host compatibility over a few generations (Woodworth *et al*. 2005; Decaestecker *et al*. 2007), and variation in host-parasite associations among adjacent island populations (Fallon *et al*. 2003, 2004, 2005; Ricklefs *et al*. 2011; Soares *et al*. 2017). Fourth, simple innate immune changes can occur in parallel between distantly related host lineages, with identical effects on susceptibility. Such a parallel change occurred in the sialic acid pathway of the ancestors of humans and owl monkeys, respectively, causing an eclectic phylogenetic distribution of susceptibility to the haemosporidian parasite, *Plasmodium falciparum* (Martin *et al*. 2005). In sum, the evidence for short-term and spatial variation in exposure and host defensive ability suggests that we should find no signal of phylogenetically conserved susceptibility across the host phylogeny.

Alternatively, we may expect phylogenetically conserved susceptibility in multi-host, multi-parasite systems either because of conserved host traits or phylogenetic constraints on parasite host range and host-switching. Many host traits that have previously been shown to affect interspecific variation in susceptibility also tend to show phylogenetic signal. These include embryonic development rates (Ricklefs 1992; Ricklefs *et al*. 2018), diet (Masello *et al*. 2018), nesting habits (Lutz *et al*. 2015), and environmental niche characteristics such as habitat and elevation (González *et al*. 2014). It remains to be adequately tested whether additional variation in susceptibility can be explained by the phylogenetic history of host species, even after taking other causes into account. Such a finding would imply that susceptibility itself is conserved, perhaps due to specific molecular genetic aspects of the host immune system, or other hidden causes. On the other hand, apparent variation in susceptibility among host species could simply be caused by variable numbers of compatible parasites, with higher prevalence expected for host species with richer parasite assemblages (Arriero & Møller 2008).

Parasite or pathogen species tend to exhibit host ranges that are phylogenetically limited, with lower infectivity, virulence, and disease intensity at increasing phylogenetic distances from the most frequently infected host species (Tinsley & Majerus 2007; de Vienne *et al*. 2009; Russell *et al*. 2009; Gilbert *et al*. 2015). This phylogenetic host-range effect has been demonstrated experimentally in fungal pathogens of plants (Gilbert & Webb 2007), rhizobial bacteria of Acacia trees (Barrett *et al*. 2016), and RNA-viruses of *Drosophila* (Longdon *et al*. 2011). Patterns of hematozoon presence-absence in New World primates also suggest a phylogenetic host-range effect (Davies & Pedersen 2008). This effect can mediate the rate and phylogenetic pattern of host-switching. The frequency of host-switching tends to be inversely proportional to the phylogenetic distance between donor and recipient host species for bat viruses (Streicker *et al*. 2010), primate lentiviruses (Charleston & Robertson 2002), plant fungal pathogens (Gilbert *et al*. 2012), and the haemosporidian-bird system that is the focus of this study (Clark & Clegg 2017). Thus, the phylogenetic distance effect appears to be a shared evolutionary feature of multi-host, multi-parasite systems.

Infection status of individuals, or infection rates of populations, provide an index of susceptibility. Some evidence of phylogenetic signal or taxonomic clade effects on infection rate has been found previously for protozoan parasites (Scheuerlein & Ricklefs 2004; Svensson-Coelho *et al*. 2013; González *et al*. 2014; Waxman *et al*. 2014; Lutz *et al*. 2015; Fecchio *et al*. 2017b), but no previous analysis has estimated phylogenetic effects while taking other relevant predictors of infection into account. This approach is essential to understanding whether susceptibility is truly conserved or simply appears conserved because shared environments or life history characteristics among related species affect infection rates. Here, we quantified phylogenetic effects on infection rate while taking into account environmental, spatial, temporal, individual, and species-level variation that could contribute to infection risk. We surveyed haemosporidians from the world’s most diverse avifauna, at the juncture of the Amazon basin and tropical Andes, including 40 host families, 523 host species, 3983 host individuals, 1678 haemosporidian infections, and 144 localities, spanning ~7 degrees of latitude, and ~4500 meters in elevation. We used phylogenetic mixed models to explicitly estimate the proportion of variance that is attributable to phylogeny and species identity, respectively. We found deeply-conserved patterns of susceptibility to haemosporidian parasites across the avian tree, demonstrating that deep phylogeny matters to real-time ecological interactions.

## MATERIAL AND METHODS

### Sampling and individual specimen data

We collected bird specimens in the Andes Mountains and adjacent lowlands (elevational range: 115–4,637 meters; Fig. 1a) in accordance with animal care guidelines and appropriate permits. Specimens and tissues are housed at the Museum of Southwestern Biology, the Field Museum of Natural History, and el Centro de Ornitología y Biodiversidad. Specimen-related data are available on the ARCTOS database (arctosdb.org) and in Table S1 of Supporting Information. Elevation, latitude, longitude, sex, body mass, and date were recorded for each specimen at the time of collection. To account for site variation in climate, we extracted 19 bioclimatic variables describing aspects of temperature and precipitation from the WorldClim database (Hijmans *et al*. 2005) based on the geographic coordinates for each specimen.

**Fig. 1.**
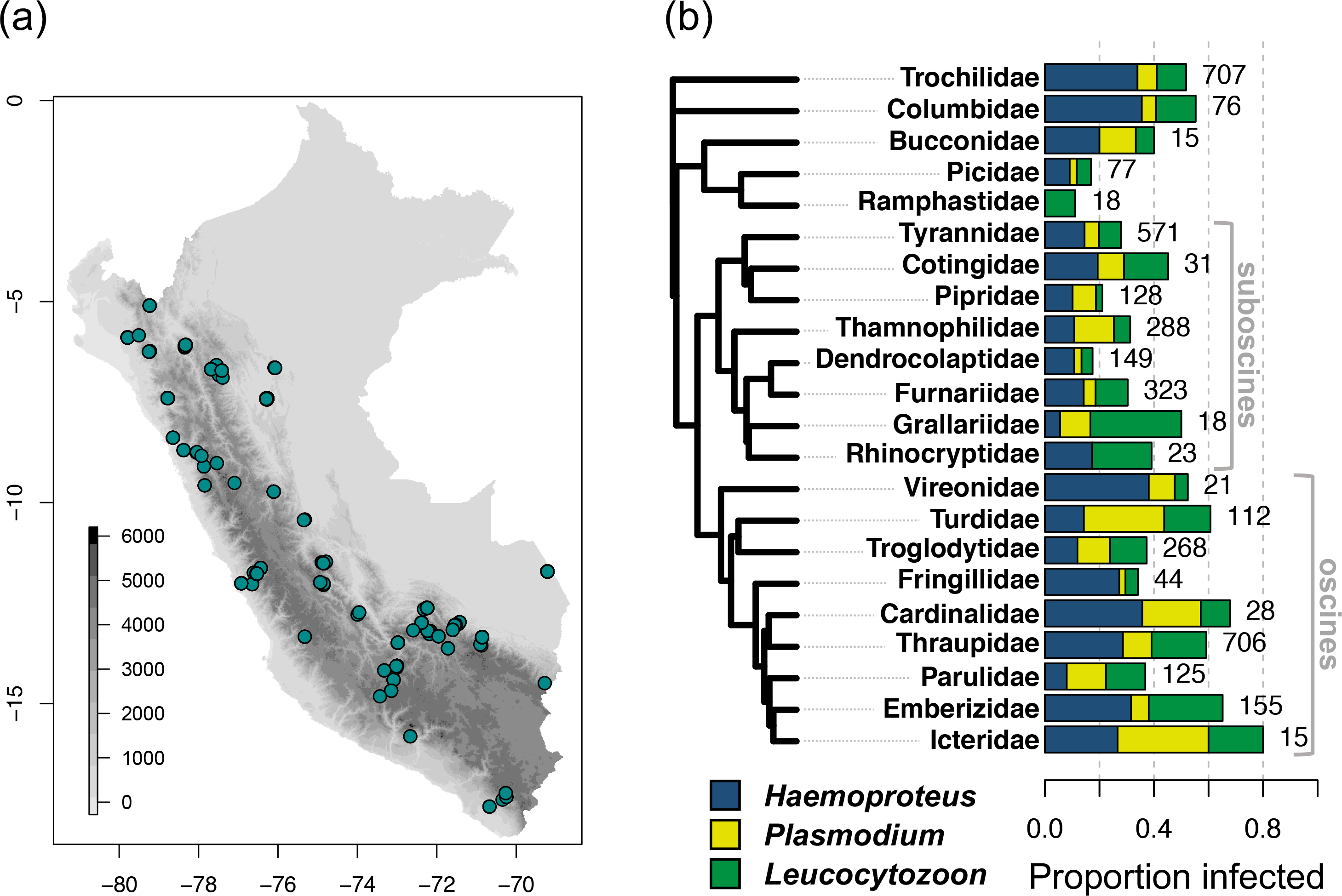
Summary of avian haemosporidian infection across Andean bird families. (a) Distribution of sample localities in Peru. Elevation is from the SRTM database with 90 m resolution, using the *raster* R package (Hijmans 2016). (b) Combined prevalence for *Haemoproteus* (including *Parahaemoproteus*; blue), *Plasmodium* (yellow), and *Leucocytozoon* (green) across well-sampled (≥15 individuals) bird families. Bar plots depicting the proportion of birds infected by each haemosporidian genus are stacked. Sample sizes are shown adjacent to bars. The tree is a least-squares consensus of 100 phylogeny subsets from BirdTree.org.

### Species-level ecological and life history traits

For each of the 523 host species, we compiled data for ecological and life history traits thought to influence haemosporidian infection status (Table S2). We obtained foraging stratum and relative abundance from the reference database published for Neotropical birds (Parker *et al*. 1996). Foraging stratum was converted to a continuous variable, with higher values indicating higher strata (1 = terrestrial, 9 = aerial). For relative abundance, we classified species into three categories: common (C), fairly common (F), or uncommon/rare (U). The remaining traits (nest type, nest height, plumage dimorphism, sociality, uniparental care, cooperative breeding, lekking, and colonial nesting/roosting) were inferred from del Hoyo *et al*. (2018) and other secondary sources. We categorized nest type as either “open” or “closed” (including cavities, domes, or nests in caves), and nest height as either “ground”, “low” (≤ 3 meters), or “high” (> 3 meters). Plumage dimorphism was classified as “none”, “moderate”, or “striking”. We categorized sociality into “solitary” (foraging alone or in pairs), “family” (family groups only), “single species” (larger groups of the same species), or “flocking” (regularly occurring in mixed species flocks). Uniparental care, cooperative breeding, lekking, and colonial nesting/roosting were classified as either “yes” or “no”. When breeding information was unavailable, we inferred the state of these traits from related species.

### Assigning infection status

For each bird, we determined infection status for all haemosporidian genera combined (overall infection), and for each genus separately (*Haemoproteus*, *Plasmodium*, and *Leucocytozoon*). We extracted DNA from tissue or blood using QIAGEN kits, and used nested PCR to amplify 478 bp of *cytb*, a mitochondrial gene (Hellgren *et al*. 2004; Waldenström *et al*. 2004; Galen & Witt 2014). PCR products were visualized on agarose gels to identify infected samples, which were then sequenced in both directions. To identify haemosporidian infections to genus, we compared them to the MalAvi database (Bensch *et al*. 2009).

### Infection across the avian tree

We used 100 trees from BirdTree.org, using the backbone tree from Hackett *et al*. (2008), that represent the range of possible phylogenetic histories among the 523 bird species in our study. Details are described in Jetz *et al*. (2012). To visualize patterns of infection across the avian tree, we first generated a consensus tree using the ls.consensus() function in the R package *phytools v. 0.6-00* (Revell 2012). For each host species, we calculated the proportion of birds infected overall and by each haemosporidian genus. We mapped infection proportions onto the tree using the contMap() function in *phytools*, which estimates the maximum likelihood ancestral states of continuous traits at internal nodes and interpolates the states along edges based on Felsenstein (1985). Estimates of prevalence have low accuracy with small sample sizes (Jovani & Tella 2006), therefore we visualized infection patterns across the phylogeny using species for which there were at least 10 samples (135 species from 17 families).

### Phylogenetic models with repeated measurements

We used principal component analysis (PCA) to summarize bioclimatic variables across the 144 unique sample localities. The first two PC axes explained 80.0% of the variation in temperature and precipitation and were included as predictor variables in models. Loadings suggested that axis 1 corresponds to increasing temperature across sites, hereafter called ‘temperature’, and axis 2 corresponds to decreasing precipitation, hereafter called ‘aridity’ (Table S3). Continuous variables (temperature, aridity, elevation, latitude, sampling month, body mass, and foraging stratum) were standardized to a mean of zero and standard deviation of one. For traits measured for multiple individuals within species, we accounted for multiple measurement effects following de Villemereuil & Nakagawa (2014). This approach uses within-group centering to separate each predictor into two components, one accounting for between-species variability and the other accounting for within-species variability. We calculated species means (between-species variability) and subtracted the mean value from individual observations (within-species differences) and included both components as predictors in models.

We built phylogenetic generalized linear multilevel models using two different Bayesian statistics packages in R: *MCMCglmm* (Hadfield 2010) and *brms* (Bürkner 2017). The packages differ in the core Bayesian algorithms they use as well as support for different model types (reviewed in Mai & Zhang 2018), but both are capable of incorporating phylogenetic information into multilevel models. We ran models using both packages and compared the outputs. *MCMCglmm* uses Markov chain Monte Carlo sampling to fit its models, whereas *brms* creates and fits models in Stan and can easily interface with the R package *loo* (Vehtari *et al*. 2017) to compute different information criteria. For both packages, we modeled bird infection as a binary response (0 for uninfected, 1 for infected) separately for each of four different outcomes: the presence of overall haemosporidian infection, and infection for each of the three genera (*Haemoproteus, Plasmodium*, and *Leucocytozoon*). For each model, predictor variables included the standardized species mean and within-species predictors for continuous individual-measured traits, as well as species-level factor predictors. These predictors encompass variation related to the environment (temperature, aridity, elevation, latitude), season (sampling month), individual (sex, body mass) or population (relative abundance) characteristics, and life history and behavior (foraging stratum, sociality, nest type, nest height, uniparental care, cooperative breeding, plumage dimorphism, lekking, and colonial nesting/roosting).

For each response, we compared ten models: 1) an intercept-only null model, 2) an intercept-only model with both species and phylogenetic random effects, 3) a model with all predictors and no random effects, 4) a model with all predictors and only a species random effect, 5) a model with all predictors and only a phylogenetic random effect 6) a model with all predictors and both phylogenetic and species random effects, 7) a reduced-predictor model with no random effects, 8) a reduced model with only a species random effect, 9) a reduced model with only a phylogenetic random effect, and 10) a reduced model with both phylogenetic and species random effects. The reduced models included only the predictors found to be important, i.e., 95% credible intervals (CI) non-overlapping with zero, in the full models. The predictors retained in the reduced models differed for each response, as reported in Results.

In *MCMCglmm*, we used the MCMCglmm() function with a “categorical” family and ran the model across four chains for 200,000 iterations with a burnin period of 100,000, thinned every 100 steps. We used default priors for the fixed effects, with priors of V = 1, nu = 0.02 for both residual and random effect variances. In *brms*, we ran models using the brm() function with the “Bernoulli” family and default priors. We ran four chains for 20,000 iterations with a burnin period of 10,000, thinned every 10 steps, for a total of 4,000 samples. For both packages, we visually checked for convergence using traceplots and confirmed that Rhat values were less than 1.01. In *brms*, we compared models using the widely applicable information criterion (WAIC, Watanabe 2010) values as well as approximate leave-one-out cross-validation information criterion based on the posterior likelihoods (LOOIC, Vehtari *et al*. 2017). We estimated fixed effects (means and 95% CI) from the posterior distributions for each predictor.

### Phylogenetic signal estimates

Phylogenetic signal, or lambda (λ), was estimated from the models as the phylogenetic heritability described by Lynch (1991). Similar to heritability in quantitative genetics, phylogenetic signal can be estimated as the proportion of the total variance attributed to the phylogenetic variance. We estimated phylogenetic signal using the full and reduced models for infection overall and for each haemosporidian genus. We also estimated the proportion of the total variance attributed to host species, which accounts for unique aspects of the susceptibility of species that are not captured by the modeled species traits, or individual or environmental characteristics. In *MCMCglmm*, the mean and 95% highest posterior density (HPD) of λ were calculated for each MCMC chain by dividing the phylogenetic variance-covariance (VCV) matrix by the sum of the phylogenetic, species, and residual VCV matrices (Hadfield & Nakagawa 2010). In *brms*, phylogenetic signal was computed following the vignette and recommendations of P. Bürkner (https://cran.rproject.org/web/packages/brms/vignettes/brms_phylogenetics.html), using the ‘hypothesis’ method and substituting π^2^/3 for the residual variance.

To assess the effect of phylogenetic uncertainty on our models and phylogenetic signal estimates, we ran the full *brms* models with 100 trees that were randomly selected from the set of most likely trees. For each tree, we estimated the inverse phylogenetic covariance matrix and ran the full *brms* model with four chains for 20,000 iterations each, including 10,000 burnin samples, thinning every 10 samples for a total of 4,000 samples. We calculated mean λ as above from the posterior distributions of each of the 100 replicate runs to determine the extent of variation in phylogenetic signal estimated under alternative phylogenetic hypotheses.

### Parasite diversity and infection rate

One additional explanation for variation in susceptibility is variation in parasite diversity; hosts that can harbor more parasite species have been shown to have higher prevalence (Arriero & Møller 2008). We tested this additional predictor of infection using the same *brms* model structure described above. To generate an estimate of parasite diversity independent of sampling and infection rate, we first pruned the host dataset to include only infected host species with at least five sequenced infections. We used the rarefy() function in the *vegan* 2.5-2 R package to produce a rarefied haplotype diversity index for each host species. This predictor was standardized as the other continuous variables described above, and included in the *brms* model with the reduced host species dataset.

## RESULTS

### Infection status summary

We detected 1,554 infected birds (39.0%), including 829 birds infected with *Haemoproteus* (20.8%), 355 with *Plasmodium* (8.9%), and 494 with *Leucocytozoon* (12.4%). Haemosporidian infection rate varied across the avian phylogeny (Fig. 1b, Fig. 2). Avian families with the highest infection rates (> 50% of birds infected) included Columbidae and several oscine Passerine families (Icteridae, Cardinalidae, Emberizidae, Turdidae, and Thraupidae). In general, we found higher rates of infection in oscines compared to suboscines, and in certain hummingbird clades (brilliants and coquettes) compared to others (emeralds and hermits) (Fig. 2; Fig. S1).

**Fig. 2.**
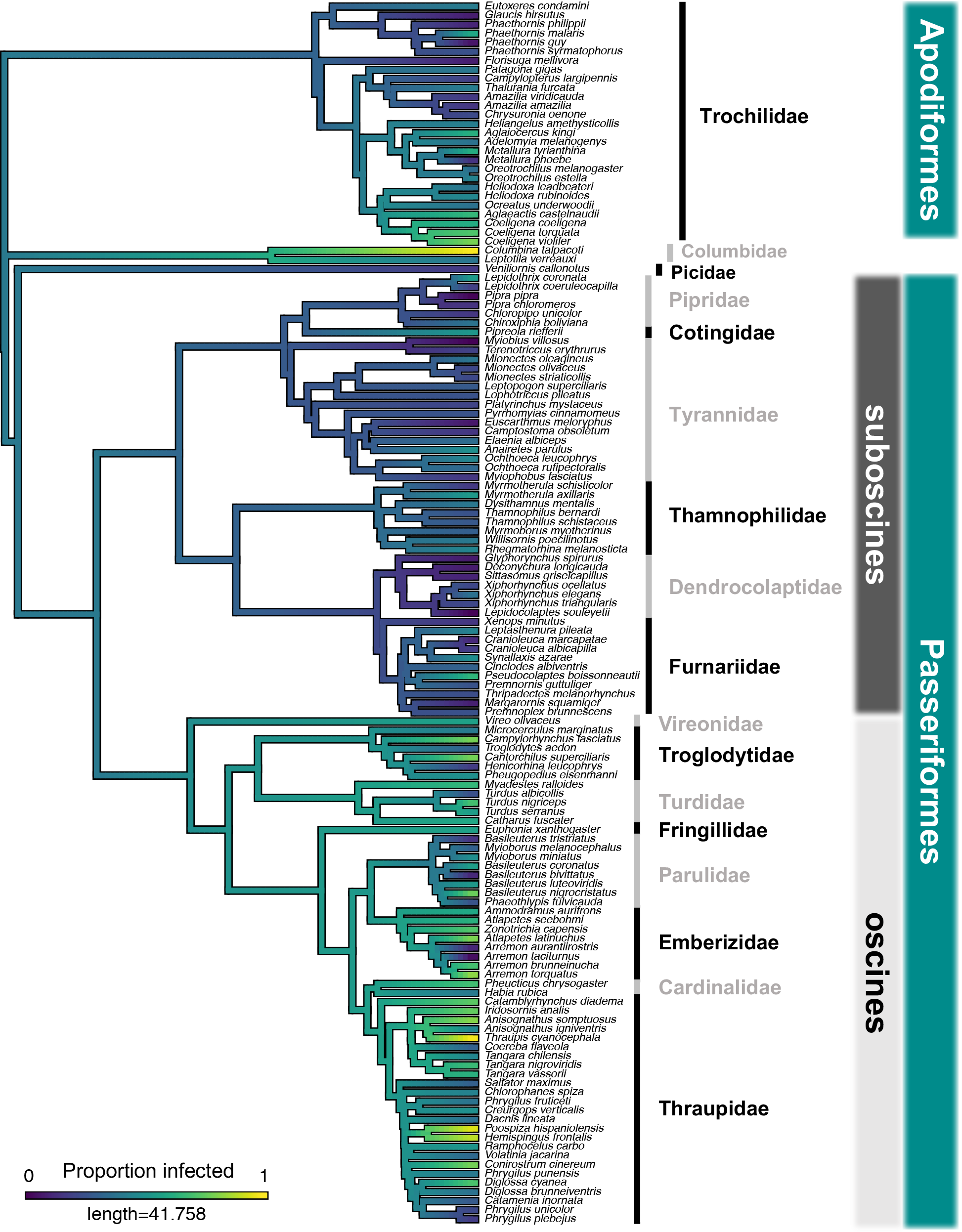
Haemosporidian infection across the avian phylogeny. The proportion of individuals infected for each well-sampled host species (≥10 individuals) was mapped as a continuous trait using the contMap() function in *phytools* (Revell 2012). The tree is a least-squares consensus of 100 phylogeny subsets from BirdTree.org.

### Predictors of haemosporidian infection

Models that included phylogenetic and species random effects fit substantially better than models with no random effects (Table 1). The reduced-predictor models with both species and phylogenetic random effects had the lowest WAIC and LOOIC scores for overall infection, *Haemoproteus*, and *Leucocytozoon*. For *Plasmodium*, the reduced-predictor model with only a phylogenetic random effect had the lowest scores. We sought to quantify the proportions of variation attributed to phylogeny and species, respectively, thus we report the results from the reduced-predictor models including both random effects for all responses.

**Table 1.** Comparison of *brms* models with species, phylogenetic (phylo), both, or no random effects. Model fit was assessed using the widely applicable information criterion (WAIC) and leave one out cross-validation (LOOIC). WAIC results were consistent with LOOIC. The difference between each model and the best fit model (lowest LOOIC) is shown as ∆LOOIC with the standard error (SE). Reduced models include the set of predictors that were considered important in the full models (95% CI non-overlapping with 0) for each response, i.e., a different set of predictors is included in each of the reduced models. For all responses, models including species and phylogenetic random effects fit substantially better than models without random effects.

| Model Description |  |  |  |  | Response |  |  |  |
| --- | --- | --- | --- | --- | --- | --- | --- | --- |
| Predictors |  |  | Random Effects |  | Overall Infected | *Haemoproteus* | *Plasmodium* | *Leucocytozoon* |
| None | All | Reduced | Species | Phylo | ΔLOOIC (SE) | ΔLOOIC (SE) | ΔLOOIC (SE) | ΔLOOIC (SE) |
| X |  |  |  |  | 521.3 (45.3) | 507.9 (46.4) | 252.2 (34.6) | 454.3 (41.3) |
|  |  | X |  |  | 434.8 (41.1) | 374.5 (38.4) | 168.4 (27.5) | 240.5 (31.7) |
|  | X |  |  |  | 335.3 (37) | 309.1 (37) | 155.4 (26.1) | 170.6 (29.8) |
|  |  | X | X |  | 36.1 (12.1) | 10.3 (7.1) | 19.5 (13.7) | 7.8 (7.2) |
|  | X |  | X |  | 33.5 (12.3) | 8.9 (10.9) | 33.2 (14.6) | 6.9 (12.3) |
| X |  |  | X | X | 22 (12.6) | 14.3 (10.1) | 28.6 (14.2) | 89.9 (18.6) |
|  | X |  |  | X | 22 (12.7) | 32.5 (15.2) | 13 (8.8) | 24.3 (15.1) |
|  |  | X |  | X | 15.2 (10.2) | 32.4 (11.7) | 0 | 20.3 (9.7) |
|  | X |  | X | X | 10.2 (7.7) | 6.4 (9.6) | 18.4 (9.7) | 2.9 (11.3) |
|  |  | X | X | X | 0 | 0 | 2.5 (4.9) | 0 |

Several predictors (foraging stratum, uniparental care, cooperative breeding, plumage dimorphism, lekking, and sociality) were unimportant for any of the responses (i.e., the 95% CI overlapped with 0) and were removed to construct the reduced models. Parameter estimates from *MCMCglmm* and *brms* were highly consistent (Fig. 3, Fig. S2–S5).

**Fig. 3.**
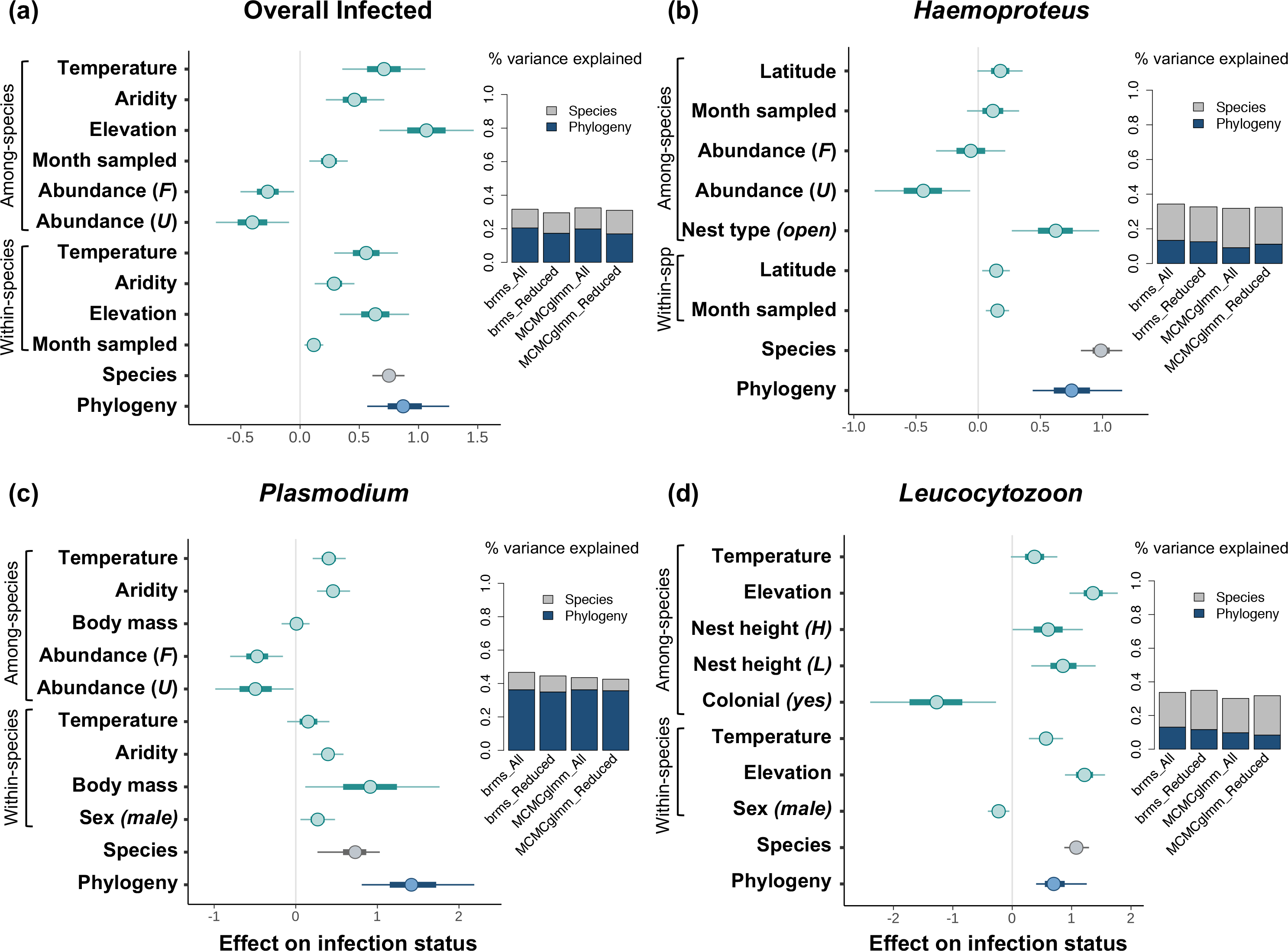
Posterior mean estimates and 95% credible intervals of predictors and random effects on infection status for reduced *brms* models. Parameters with intervals that do not overlap zero are considered to have a significant influence on the response. Intercepts were removed for visualization and are shown in Fig. S4. For continuous variables, both among-species and within-species effects are shown. For categorical variables, the effects shown are relative to the reference categories: Sex *(female)*, Nest type *(closed)*, Abundance *(common)*, Nest height *(ground)*, and Colonial *(no)*. F = Fairly common, U = Uncommon/rare, H = high, L = low. Panels to the right depict the proportion of total variance attributed to species (grey) or phylogeny (blue). Results are consistent across full and reduced models for both *MCMCglmm* and *brms*. The proportion of variance attributed to phylogeny is the phylogenetic signal, which was significant for all models based on the ‘hypothesis’ test in *brms*.

Aspects of climate, elevation, and latitude were important for overall infection and for each haemosporidian genus. Overall infection probability increased with increasing temperature, aridity, elevation, and sampling month (Fig. 3a). Host species that were less abundant (fairly common or uncommon) tended to be less infected than common species. Different species-level predictors were considered important for susceptibility to each haemosporidian genus (Fig. 3).

*Haemoproteus* infection increased slightly with latitude and sampling month and was lower for uncommon host species compared to common species (Fig. 3b). Species with open nests tended to have higher *Haemoproteus* infection compared to species with closed nests.

*Plasmodium* infection increased with temperature, aridity, and within-species body mass (Fig. 3c). Male hosts had higher *Plasmodium* infection compared to females, and species with lower abundances (fairly common or uncommon) tended to be less infected than common species.

*Leucocytozoon* infection increased with increasing temperature and elevation (Fig. 3d). *Leucocytozoon* infection was lower for males than females, higher for species with low nests (<3 m) than those with ground nests, and lower for colonial species than non-colonial species.

### Phylogenetic signal in infection

Phylogenetic signal was important for all models; 95% CI of phylogenetic random effects do not overlap with 0 (Fig 3). The proportions of total variance attributed to phylogeny and species, respectively, were consistent between full and reduced models for both *MCMCglmm* and *brms* (Table 2, Fig. 3). For the reduced *brms* models shown in Fig. 3, phylogenetic signal in overall infection was 0.17 [95% CI 0.06–0.33]. Phylogenetic signal was lower for *Haemoproteus* (0.13; [95% CI 0.03–0.28]) and *Leucocytozoon* (0.12; [0.03–0.3]), and highest for *Plasmodium* (0.35; [0.11–0.61]). For *Haemoproteus* and *Leucocytozoon*, the variance attributed to host species was larger than the variance attributed to phylogeny (Fig. 3). Our phylogenetic signal estimates were consistent under alternative, plausible phylogenetic hypotheses, indicating that the results are robust to phylogenetic uncertainty (Fig. S6). Running the full *brms* model with 100 trees randomly selected from the set of most likely trees produced a mean λ = 0.24 (range: 0.18–0.31).

**Table 2.**
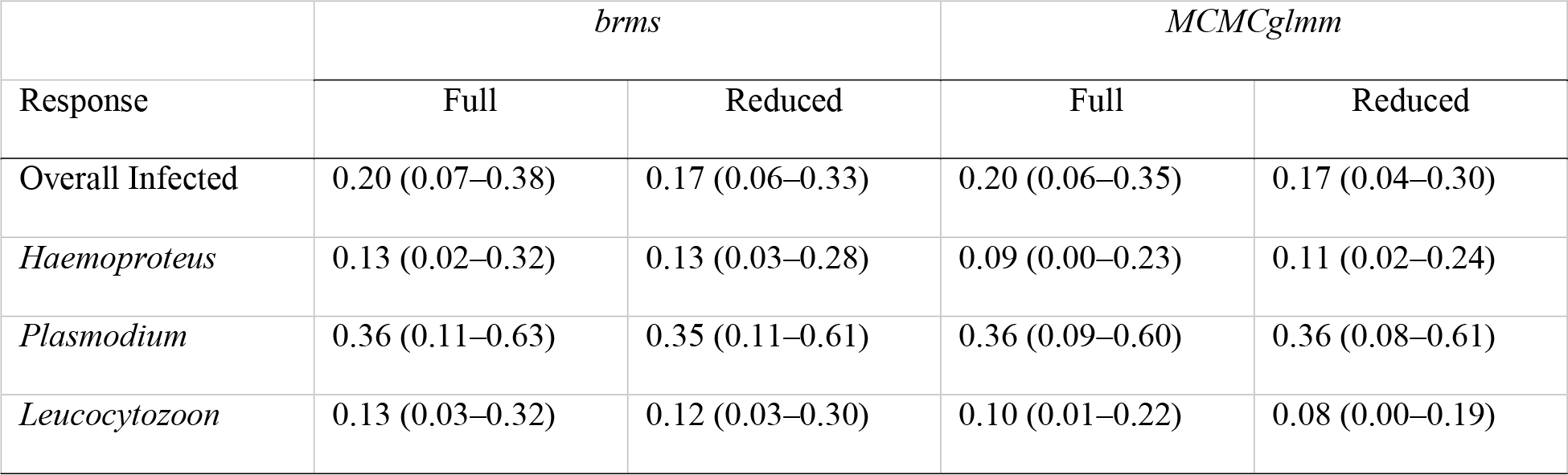
Phylogenetic signal estimates from *brms* and *MCMCglmm* full and reduced models. Means and 95% credible intervals for phylogenetic signal, or lambda (λ), estimated from *brms* and *MCMCglmm* for full and reduced models. λ is the proportion of total variance attributed to phylogenetic variance.

### Parasite diversity and infection rate

Of the 367 infected species, 112 included five or more sequenced infections (mean = 11, max = 49 infections). Within host species, rarefied haplotype diversity ranged from one to five (mean = 3.98, stdev = 0.93). Model results indicated that parasite diversity was not an important predictor of overall infection rate (Fig. S7).

## DISCUSSION

Variation in susceptibility among host species is a common feature of multi-host, multi-parasite systems, but the importance of phylogeny, while taking other predictors into account, has rarely been addressed. We found that host phylogeny explains substantial variation in haemosporidian infection rate, indicating that susceptibility is conserved on the time scale of avian diversification. Our statistical approach accounted for a suite of life history, behavioral, temporal, and environmental effects that are demonstrated drivers of infection rate. The results were consistent between two Bayesian modeling methods and with reduced sets of predictors. Phylogenetic and species effects were important for all parasite genera, but differed in magnitude of effect. Phylogenetically conserved susceptibility should affect many aspects of the evolutionary dynamics of multi-host, multi-parasite systems, including biogeography, ecoclimatic niches, diversification rates, and host-switching patterns.

Phylogenetic conservation of susceptibility was evident at remarkably deep levels within the avian tree. Most notably, oscine songbirds exhibited substantially higher infection rates than their sister clade, the suboscines (Fig. 1, Fig. 2). These two clades account for most of the diversity on the ‘bird continent’, and are known to differ in fundamental ways, including sound production mechanisms, song learning, pigmentation, and metabolic rate (Kroodsma 1983; Swanson & Bozinovic 2011). The current study confirms that they also differ with respect to susceptibility to haemosporidian parasites, with suboscines being consistently less infected (Ricklefs 1992, 2002).

Some environmental characteristics clearly influence infection rates, while others do not. For example, overall infection rate tended to increase with increasing temperature and aridity, and *Leucocytozoon* infection increased substantially at higher elevations. This finding is consistent with previous studies demonstrating different elevational patterns of infection rate among haemosporidian genera, possibly caused by elevational variation in vector abundance and exposure rate (van Rooyen *et al*. 2013; González *et al*. 2014; Harrigan *et al*. 2014).

Life history and ecological factors also explain some variation in infection among species. For example, *Haemoproteus* infection was higher for species with open nests compared to closed nests, and *Leucocytozoon* infection was higher for species with midstory (<3 m) nest heights compared to ground nesters. Results from previous studies that have addressed these factors are mixed (Svensson-Coelho *et al*. 2013; Lutz *et al*. 2015; Fecchio *et al*. 2017a), with effects typically attributed to vector ecology. Vector feeding preferences for certain host species affects susceptibility, at least within depauperate communities (Medeiros *et al*. 2015); nevertheless, compatibility of haemosporidian parasites with host species immune systems remains an overarching determinant of host susceptibility (Medeiros *et al*. 2013).

Phylogenetic variation in susceptibility rises above the variation explained by environmental and species traits, including a suite of traits that should explain variation in exposure. This is remarkable for at least two reasons. First, an evolutionarily labile feature such as an ecological interaction that fluctuates in real time would not be expected to remain predictable on the basis of distant phylogenetic affinities. Second, several of the environmental and life history traits that explain some variation in susceptibility are themselves subject to phylogenetic signal; it is striking that there is additional phylogenetic signal even after these conserved predictors are included in the model. The conserved evolution of infection status is a distinct and interesting aspect of phylogenetic niche conservatism (Wiens *et al*. 2010), wherein ecological interactions are sustained long-term during divergence of related lineages.

The causes of deep phylogenetic conservation of susceptibility are most likely molecular genetic aspects of the innate immune system that are also phylogenetically conserved, with specificity that is lost gradually over evolutionary time (Schulze-Lefert & Panstruga 2011). Many innate immune factors are deeply conserved and subject to strong purifying selection (Malo *et al*. 1994; Hückelhoven *et al*. 2013), but disease resistance can also evolve through single substitutions in these factors, often accompanied by negative pleiotropy (Aidoo *et al*. 2002; Carter & Nguyen 2011). The latter mechanism could explain the species random effects demonstrated by our analyses. The basis of species-specificity in malaria parasites can be as simple as a single, large effect mutation, such as the single change to the CMAH gene in the ancestor of *Homo sapiens* that became the basis for host specificity in *Plasmodium falciparum* and related *P. reichenowi* in chimps (Martin *et al*. 2005).

The closest precedent for our finding is that of Longdon *et al*. (2011), who found phylogenetic signal in viral susceptibility among *Drosophila* species that were experimentally infected with sigma viruses. In that case, susceptibility of host species to each of three sigma viruses tested was correlated, indicating that it resulted from variation in the generalized immune response. In our study, infection rates for the three haemosporidian genera within host families and host species were largely uncorrelated (Fig. S8). This lack of correlation may be a result of specialized ecoclimatic niches of haemosporidian genera and host lineages, respectively, causing general susceptibility to manifest differently in different environments. We found that phylogenetic signal in susceptibility was strongest in *Plasmodium*, the most host-generalized of the three genera (Valkiu☐nas 2005). Accordingly, we suggest that our results, like those of Longdon *et al*. (2011), are consistent with variation in the generalized immune response.

The success of a parasite depends on the interaction between host resistance traits and parasite counter-adaptations. Classical defense theory holds that faster growth of the host will confer lower resistance to parasites (García-Guzmán & Heil 2014). Increased resistance could thus be explained by slower host-development time, as has been suggested in grass-virus (Cronin *et al*. 2014), amphibian-trematode (Johnson *et al*. 2012), and bird-haemosporidian systems (Ricklefs 1992; Ricklefs *et al*. 2018). Variation in host development rate could be a latent variable that contributed to the phylogenetic signal in susceptibility that we observed.

Abundance of close relatives in a host community may also affect infection rate (Parker *et al*. 2015; Ellis *et al*. 2017), and could be conserved across our sample of host species. The lack of a relationship between susceptibility and species diversity of host families in the Peru avifauna suggests this did not impact our results (Fig. S9). We also included relative abundance in our models and found that although more abundant species did tend to be more infected, phylogeny still explained substantial variation in infection status.

If conserved host ranges are a general tendency of parasite clades (Gilbert & Webb 2007; Davies & Pedersen 2008; de Vienne *et al*. 2009; Russell *et al*. 2009; Longdon *et al*. 2011), it could cause related host species to harbor similar parasite communities. Such a process could plausibly lead to phylogenetic signal in infection rates in two ways. First, if certain host clades diversified more rapidly, species within those clades may receive host switches at higher frequency and exhibit higher susceptibility. Indeed, Engelstädter & Fortuna (2018) predicted higher infection rates in faster diversifying clades as a consequence of host-shifts tending to be among close relatives. We found no clear evidence of this pattern in our data; host family-level diversity, whether global or regional, was not linked to infection rate (Fig. S9). Secondly, the phylogenetic host-range effect could result in a conserved pattern of susceptibility if particular host clades have more compatible parasites than others. In this case, one prediction is that host species within clades that have higher parasite diversity would have higher susceptibility. In this study, we found no evidence for an effect of parasite diversity on infection rate (Fig. S7). Furthermore, for avian haemosporidians, the community of parasite lineages in any given host species or family tends to be drawn from across the haemosporidian phylogeny (see Fig. 1b), and generalist parasites with eclectic host ranges are frequent (Hellgren *et al*. 2009; Loiseau *et al*. 2012; Svensson-Coelho *et al*. 2013). This suggests that phylogenetic signal in susceptibility is not explained by conserved host ranges in this system.

Phylogenetic conservatism of species interactions could have implications for evolutionary fates and net diversification rates of clades. The results of this study underscore how deep evolutionary history is relevant to real-time ecological outcomes, suggesting there are constraints on immune system innovations that affect long-term shifts in susceptibility. However, the implications for diversification are not simple; we found no link between infection rate and host-clade size at the family level (Fig. S9). Phylogenetic variation in susceptibility implies that changes in disease pressure are likely to affect the phylogenetic structure of communities (and vice versa) and could potentially maintain phylogenetic alpha- and beta-diversity (Barrett *et al*. 2009). Alpha-diversity could be enhanced via Janzen–Connell type dynamics (Terborgh 2012; Gilbert & Parker 2016), in which density of conspecific or related hosts is regulated by shared susceptibility to a parasite. Beta-diversity (and alpha-diversity) could be enhanced by ‘apparent competition’ (Holt 1977; Ricklefs 2010), in which species are differentially susceptible to a shared generalist parasite. The fact that some variation in susceptibility is conserved on a scale of tens of millions of years suggests that these same ecological mechanisms maintain deep phylogenetic diversity of hosts, a long-celebrated characteristic of the South American avifauna.

## ACKNOWLEDGEMENTS

We thank John M. Bates, Shannon Hackett, Emil Bautista, Shane G. DuBay, Ariel M. Gaffney, C. Jonathan Schmitt, Andrew B. Johnson, Laura Pages Barcelo, and Ben Winger. This work was supported in part by NSF DEB-1146491, NSF DEB-1503804, NSF PRFB-1611710, a CETI seed grant (NCRR-NIH P20RR018754), the Davee Foundation, the Faulk Medical Research Trust, and the Pritzker DNA Laboratory.

